# Operant training for highly palatable food alters translating mRNA in nucleus accumbens D2 neurons and reveals a modulatory role of *Neurochondrin*

**DOI:** 10.1101/2023.03.07.531496

**Authors:** Enrica Montalban, Albert Giralt, Lieng Taing, Yuki Nakamura, Assunta Pelosi, Mallory Brown, Benoit de Pins, Emmanuel Valjent, Miquel Martin, Angus C. Nairn, Paul Greengard, Marc Flajolet, Denis Hervé, Nicolas Gambardella, Jean-Pierre Roussarie, Jean-Antoine Girault

## Abstract

**BACKGROUND:** Highly palatable food triggers behavioral alterations reminiscent of those induced by addictive drugs. These effects involve the reward system and dopamine neurons, which modulate neurons in the nucleus accumbens (NAc). The molecular mechanisms underlying the effects of highly palatable food on feeding behavior are poorly understood.

**METHODS:** We studied the effects of 2-week operant conditioning of mice with standard or isocaloric highly palatable food. We investigated the behavioral effects and dendritic spine modifications in the NAc. We compared the translating mRNA in NAc neurons identified by the type of dopamine receptors they express, depending on the type of food and training. We tested the consequences of invalidation of an abundant downregulated gene, Ncdn (Neurochondrin).

**RESULTS:** Operant conditioning for highly palatable food increases motivation for food even in well-fed mice. In control mice, free access to regular or highly palatable food results in increased weight as compared to regular food only. Highly palatable food increases spine density in the NAc. In animals trained for highly palatable food, translating mRNAs are modified in NAc dopamine D2-receptor-expressing neurons, mostly corresponding to striatal projection neurons, but not in those expressing D1-receptors. Knock-out of Ncdn, an abundant down-regulated gene, opposes the conditioning-induced changes in satiety-sensitive feeding behavior and apparent motivation for highly palatable food, suggesting down-regulation may be a compensatory mechanism.

**CONCLUSIONS:** Our results emphasize the importance of mRNA alterations D2 striatal projection neurons in the NAc in the behavioral consequences of highly palatable food conditioning and suggest a modulatory contribution of Ncdn downregulation.

## INTRODUCTION

Food palatability is a potent drive to eat even in the absence of actual caloric need. Excessive consumption of highly palatable food can disrupt the normal regulation of appetite (1), induce the development of compulsive-like approach to food intake, and ultimately lead to overweight and obesity (2-6). However, the mechanisms by which exposure to palatable food induces persistent behavioral alterations responsible for maladaptive food consumption remain poorly understood.

Feeding behavior depends not only on homeostatic and metabolic regulations, but also on reward processing mechanisms including the incentive and seeking aspects of feeding. One hypothesis is that exposure to palatable food recruits the brain reward system, switching food-seeking behavior from flexible, goal-directed actions, to inflexible, compulsive-like responses (7-10). A large body of evidence supports impaired flexibility in obese subjects (11), decreased sensitivity to satiety-mediated devaluation of food (12, 13), and stronger devaluation of delayed rewards (14). In obese subjects these behavioral impairments are accompanied by alterations within the brain reward system. Functional imaging of the fronto-striatal circuit, a main substrate for inhibitory behaviors and cognitive control, reveals blunted activation in response to food and associated cues (5, 15, 16). The functionality of the fronto-striatal circuit depends on dopamine (DA) transmission and evidence supports a role of DA as a sensor of peripheral metabolic signals (17) and in mediating the value of food-associated cues (18). Downstream of DA, striatal D2 receptors (D2R, usually not distinguished from D3R) have attracted a lot of attention since D2R antagonists induce weight gain (19-22). Decrease (23), increase in ventral regions (24), or no change (25) in D2R availability have been reported in the striatum of morbidly obese subjects. Moreover, polymorphisms in ANKK1/DRD2 (DRD2 being the D2R gene) were associated with blunted signal in fronto-striatal circuits in response to food stimuli, in relation with weight gain (26).

The nucleus accumbens (NAc) is a key region in the response to both natural (food) and non-natural (drugs of abuse) rewards (27, 28). The NAc also mediates food motivation and decision-making in obesity (29). The NAc comprises two populations of projection neurons, D1R-or D2R-expressing striatal projection neurons (SPNs), also known as medium-size spiny neurons, referred to below as D1SPNs and D2SPNs, respectively (30). Early optogenetic studies indicated that D1SPNs and D2SPNs have opposing effects on reinforcement (31, 32), but further work showed that in the NAc both populations can encode reward and aversion depending on stimuli and activity patterns (33).

In animal models palatable – usually highly caloric – food exposure alters reward-seeking behaviors (34-36) with increased impulsivity (37) and impaired cognitive functions (38), which are at least partially independent of overweight. These behavioral dysfunctions are accompanied by modifications of DA transmission markers, with a decrease in D2R expression in the striatum (2, 39). However, the amplitude, direction and nature of these alterations differ between studies, possibly due to the composition of the palatable food, and duration and age of exposure (40). Yet, consistent findings point to alterations in excitatory transmission and structural plasticity of glutamatergic synapses in the striatum (41-43), further supporting the central role of this structure in palatable food-induced long-term behavioral adaptations. However, the molecular bases of these regulations are poorly understood and the impact of palatable food exposure on the transcriptional landscape of striatal neurons is not known. In many studies the effects of palatability per se are not distinguished from the metabolic and neurobiological modifications related to (over)consumption of the food. Moreover, the passive – or forced – access to the highly palatable food in rodent models does not allow isolating the specific impact of palatability on the goal-directed component of food-seeking behavior that is crucial to the development of compulsive eating.

Here, we used a combination of behavioral and genome-wide approaches to characterize alterations in the translatome associated with food-seeking behavior for standard or isocaloric highly palatable food in the main dopaminoceptive neuronal populations of the NAc. We investigated changes in D1SPNs and D2SPNs using translating ribosome affinity purification (TRAP) (44, 45) followed by RNAseq (TRAP-seq) in mice (46). We compared mice that learned to nose-poke to obtain either regular food or highly palatable food to yoked controls, which received the same food non-contingently. After identifying changes in translating mRNA in D2SPNs, we tested the consequences of the genetic manipulation of one of the genes differentially regulated by conditioning for highly palatable food, Ncdn (coding for neurochondrin, a.k.a. norbin).

## Materials and Methods

### Animals

For translatome analyses we used male and females (10-12 weeks) *Drd1*-EGFP/Rpl10a or *Drd2*-EGFP/Rpl10a mice (45), maintained as heterozygotes on a C57BL/6J background (**Supplementary Table 1**). To generate Ncdn-cKO mice, *Ncdn*^F/F^ mice were crossed with *Ncdn*^F/+^ / *Camk2a*-Cre^*/+^ double mutant mice (47). Male and female littermates were used at 3-4 months of age. C57BL/6 (WT) male mice were purchased from Janvier lab and used at 3-4 months. Animal protocols were performed in accordance with the National institutes of Health *Guide for the Care and Use of Laboratory Animals* and approved by The Rockefeller University’s Institutional Animal Care and Use Committee (#IACUC 14753-H).

### Behavioral experiments

*Operant conditioning experiments* were carried out in Med Associate operant chambers (model ENV-307A-CT) with 2 nose-poke holes (see **Supplementary Methods** for details). Mice were randomly assigned to one of the following 4 groups, differing by their role and the food they received: “master highly palatable” (mHP), “master standard” (mST), “yoked highly palatable” (yHP), “yoked standard” (yST) (**Figure 1A**). Five days before the start of conditioning all the mice were food-restricted to maintain their weight stable, until the 9^th^ operant training session in order to facilitate the acquisition of the task, and then had ad libitum food access. During the operant conditioning sessions animals were presented with either 20 mg dustless precision ST pellets (TestDiet 5UTM #1811143) or HP isocaloric pellets with a higher level of sucrose among carbohydrates and a chocolate flavor(TestDiet 5UTL #1811223). The progressive ratio (PR) schedule lasted for 1 h (see **Supplementary Methods)**. *For the free choice paradigm*, mice were separated in two groups. In their home cage, one group had only access to standard food, while the other group had free access to highly palatable and standard food. Their weight was monitored.

**Figure 1:**
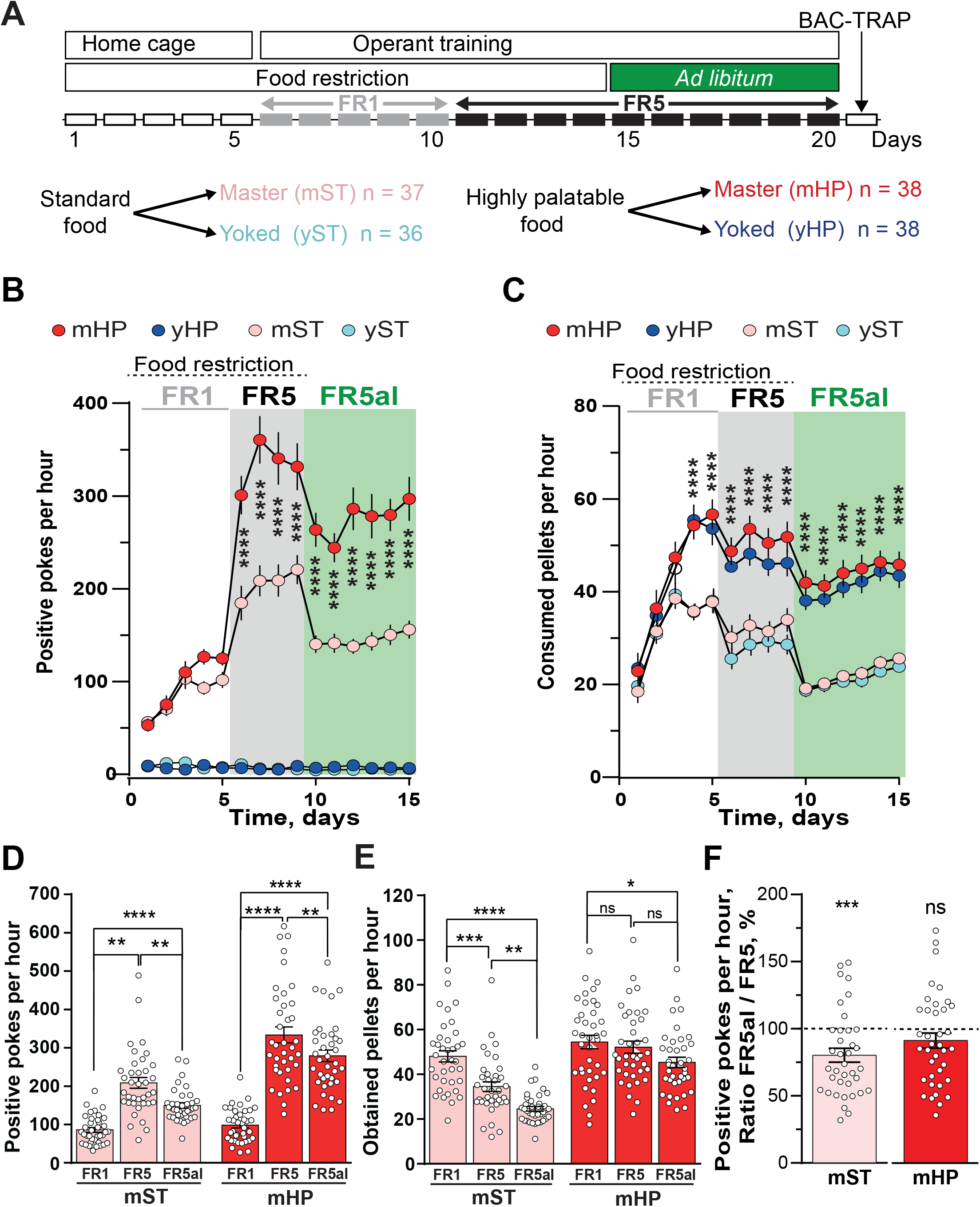
Operant training for standard and highly palatable food results in different behavioral changes. **A**) Schematic description of the operant training protocol used in this study. FR, fixed ratio. **B**) Time-course of the number of positive pokes (i.e. in the active hole) during training as described in A). Two-way repeated measure ANOVA for each phase (see **Supplementary Table 2** for all statistical results). Group effect, FR1, F_(3, 141)_ = 97.3, p<10^−4^, FR5, F _(3, 144)_ = 161, p<10^−4^, FR5al, F_(3, 144)_ = 234.5, p<10^−4^. Holm-Sidak’s multiple comparisons are indicated between mHP and mST. As expected the yoked mice did not make positive pokes. **C**) Time-course of consumed pellets. Two way repeated measure ANOVA, group effect, FR1, F_(3, 142)_ = 8.25, p<10^−4^, FR5, F_(3, 142)_ = 23.9, p<10^−4^, FR5al, F_(3, 142)_ = 50.1, p<10^−4^. Holm-Sidak’s multiple comparisons are indicated between mHP and mST. The yoked mice consumed virtually the same amount of food as their respective masters. **D**) Summary of positive pokes per hour during the 3 phases for mHP and mST as in Fig. 1B (same data). Two-way ANOVA, food effect, F_(1, 219)_ = 70.9, p<10^−4^, training effect, F_(2, 219)_ = 99.8, p<10^−4^, interaction, F_(2, 219)_ = _(1, 219) (2, 219) (2, 219)_ 13.2, p<10^−4^, Holm-Sidak’s multiple comparisons as indicated. **E**) Summary of obtained pellets per hour during the 3 phases for mHP and mST. Two-way ANOVA, food type, F = 59.4, p<10^−4^, training phase, F = 23.5, p<10^−4^, interaction, F = 5.0, p = 0.075, Holm-Sidak’s multiple comparisons as indicated. **F**) The satiety-induced decreased motivation for nose-poking was evaluated by calculating for each mouse the ratio of the average number of positive pokes per day during the **FR5al** sessions divided by the average number of positive pokes per day during the **FR5** sessions with food restriction. Data are plotted for mST (**left panel**, n = 37) and mHP (**right panel**, n = 38). One sample t test, mST, t_36_ = 3.8, p = 0.0005, mHP, t_37_ = 1.55, p = 0.13, ns. **B-F**, * p<0.05, ** p<0.01, *** p<0.001, **** p<10^−4^.

### Cell population-specific mRNA immunoprecipitation, libraries, and sequencing

Cell population-specific purification of translating mRNA was performed as described (44, 45) with some modifications (46). The brain was rapidly dissected and sliced on ice. Bilateral pieces punched out from NAc of 3 mice (**Supplementary Table 1**) were homogenized in ice-cold lysis buffer (see Ref (46) and **Supplementary Methods**). The quantity of obtained RNA was determined by fluorimetry using Quant-iT Ribogreen, and quality was checked using a Bio-Analyzer Pico RNA kit before library preparation. Five ng of RNA were used for reverse-transcription, ultra-sonicated and library construction with an Illumina TruSeq RNA sample prep kit. The libraries were sequenced on an Illumina HiSeq 2500 instrument. Details are available in **Supplementary Methods**.

### Bioinformatic analysis

The quality of the raw data was assessed using FastQC (48) and the libraries (37-62 million 50-bp paired-end reads) mapped to the *Mus musculus* genome GRCm38 (UCSC mm10) using HISAT2 (49). Reads were quantified using the RNA-Seq pipeline of SeqMonk (50). Sequencing data are deposited in NCBI’s Gene Expression Omnibus (51) (GEO # GSE137153 https://www.ncbi.nlm.nih.gov/geo/query/acc.cgi?acc=GSE137153) (**Supplementary Table 1**). Differentially expressed genes were identified with R using the Bioconductor package DESeq2 v1.30.1 (52). Genes with adjusted p-value <0.05 with the false discovery rate method (53) were declared differentially expressed. Gene ontology (GO) over-representation analyses were performed http://geneontology.org/ with GO Ontology database DOI: 10.5281/zenodo.6799722 Released 2022-07-01. Network inference was performed with combined results of CLR (54) and GENIE3 (55) as described (46), and visualized and analyzed using Cytoscape (56). We retained only the 1 % highest-ranking non-zero edges (313,944 edges). We filtered the list of edges to retain those linking genes differentially expressed between master and yoked animal fed with HP food (**Supplementary Methods**).

### Spine analysis

Spines were stained using the Golgi-Cox method (57) and counted as described in the **Supplementary Methods** by investigators blind to the mouse group.

### Statistical analyses

Data were analyzed with GraphPad Prism-6. Normal distribution was checked with D’Agostino … Pearson normality test. If n was <7 or distribution significantly different from normal, non-parametric tests were used. Complete statistical analyses results are presented in **Supplementary Table 2**.

## RESULTS

### Food palatability induces differential behavioral responses in an operant training paradigm

The mice used for translatome profiling were trained in an operant paradigm for obtaining either standard food (mST) or highly palatable food (mHP, **Figure 1A**). Yoked control mice (yST and yHP, respectively) were placed in the same conditions except that they received food pellets non-contingently when their paired master mouse obtained one. Under food restriction and low operant schedule (fixed ratio 1, FR1, i.e. one pellet obtained for one poke) mHP mice displayed slightly more positive pokes than mST animals (**Figure 1B**, for statistical analyses see **Supplementary Table 2**). As expected, yoked mice exerted no operant responses (**Figure 1B**), although they consumed the same number of pellets as their respective masters (**Figure 1C**). The increase in ratio requirement (FR5) led to a higher augmentation in operant behavior in the mHP group than in mST (**Figure 1B, D**). The number of obtained rewards did not decrease in mHP as compared to FR1 (**Figure 1E**). These data suggest that the motivational drive was enhanced in the mHP group, in accordance with the incentive effect of palatability on operant responding (58). When mice were no longer food-restricted in their home cage (*ad libitum* condition), both groups tended to decrease the number of pokes (**Figure 1B, D**). However, this decrease was more pronounced in the mST group than in the mHP group (**Figure 1D, F**), suggesting that mHP animals are less sensitive to satiety-induced devaluation of the food. These results demonstrate that, independently of the caloric intake, HP food enhances the motivational drive to seek food and blunts the effect of satiety on food-seeking behavior, two characteristics that are believed to be main culprits for the development of compulsive eating and obesity (1).

To determine whether HP food used for operant training induced a loss of control over food intake and weight gain, we evaluated the spontaneous behavior of different groups of wild type mice, which had either only access to ST food or free choice between these ST and HP isocaloric food for 30 days (**Figure 2A**). Exposure to ST/HP free choice led to a more pronounced weight gain than free access to ST food only. Together these results indicate that our groups of mice behaved as expected with a higher incentive drive for HP food.

**Figure 2:**
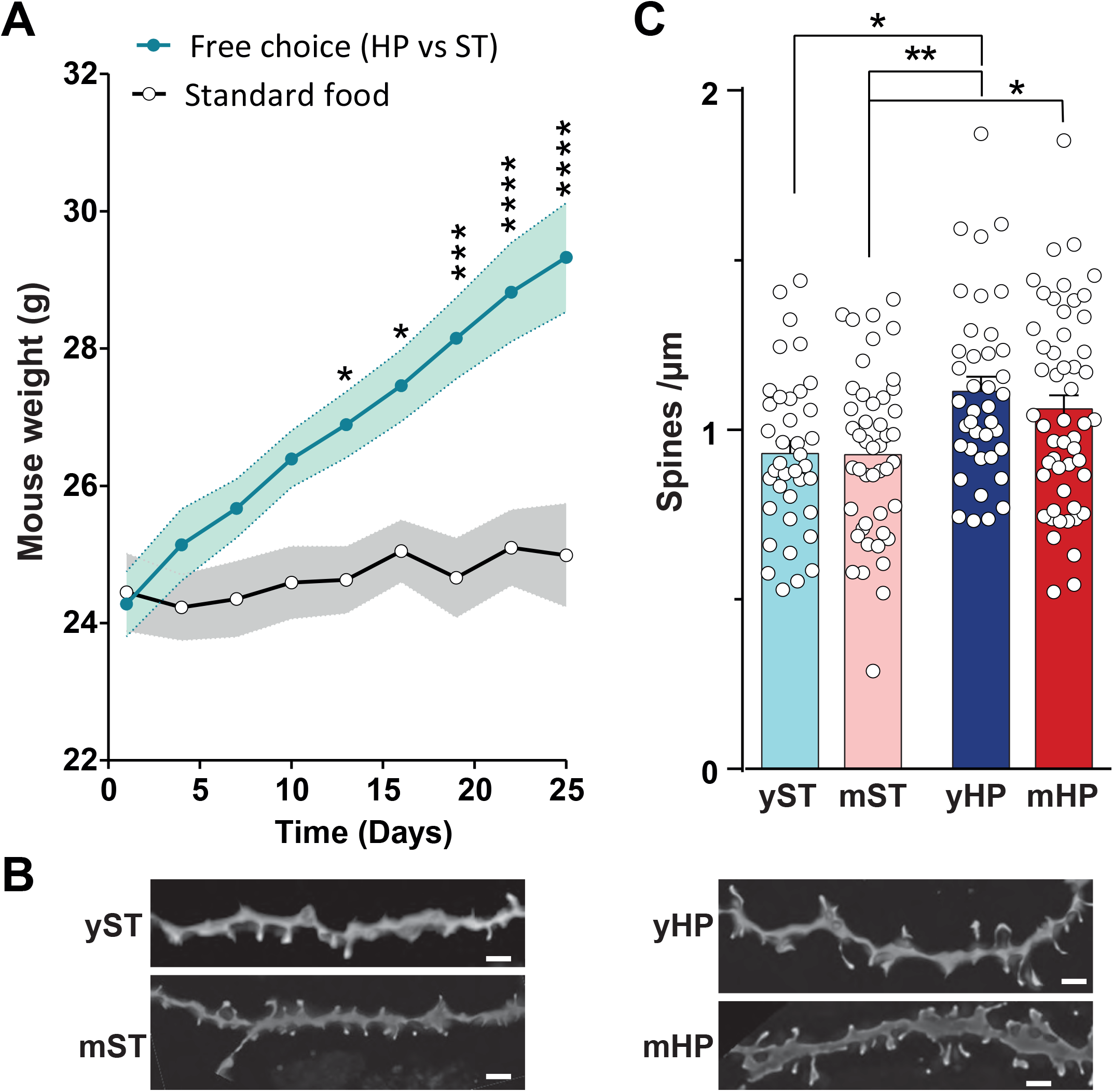
In wild type mice HP food induces a weight gain in free access conditions and increases spine density in the NAc. **A**) Male 3-4 month-old C57BL/6 mice were kept in their home cage with either free access to both HP and ST isocaloric food (Free choice, n = 10). A different group had only access to ST food (n = 10). The mouse weight was monitored every 3^rd^ day during 24 days. The mice in free choice conditions gained more weight than those with access to ST food only. For each time point, within each group the data distribution was not different from normal. Two-way repeated measure ANOVA: Interaction, F _(8,144)_ = 14.18, p <10^−4^, time effect, F _(8,144)_ = 26.30, p <10^−4^, food type effect, F_(1,18)_ = 10.09, p = 0.0052. Multiple comparisons Holmes-Sidak’s test, * p<0.05, *** p<0.001, **** p <10^−4^. **B**) Wild type mice as in A) were subjected to operant training as described in Fig. 1A. Mice were killed 24 h after the last session and sections of the NAc were stained using the Golgi-Cox methods. Examples of dendrites from mice in each group are shown. Scale bar, 4 µm. **G**) Wild-type mice (n = 8 group) were trained as in **A**), and sections through the nucleus accumbens stained with the Golgi method and the number of dendritic spines per µm counted (38-52 dendrites counted per group), two-way ANOVA, interaction F _(1, 171)_ = 0.40, p = 0.53, food type, F_(1, 71)_ = 15.90, p<10^−4^, role (yoked vs. master), F_(1, 171)_ = 0.47, p = 0.49. Imaging (B) and spine analysis (C) were done by investigators blind to the group. (**A and C**) Multiple comparisons Holmes-Sidak’s test, * p<0.05, ** p<0.01, *** p<0.001, **** p <10^−4^.

The instauration and maintenance of goal-directed behavior have been related to morphological alterations in the cortico-striatal pathway with an increase in spine density following operant training for drugs (59) or HP food (60). We therefore examined whether the exposure to HP food and/or the operant training had measurable effects on NAc neuron spine density in our experimental conditions, using Golgi staining in wild type C57BL/6 mice (**Figure 2B**). Using 2-way ANOVA, we found a food type effect (F_(1, 171)_ = 15.90, P < 10^−4^, **Figure 2C, Supplementary Table 2**). These results indicated a predominant effect of food palatability on NAc neuron spines with an increased density after 2-week training.

### TRAP-seq reveals major effects of palatable food conditioning on gene expression in D2SPNs

Since long-lasting behavioral adaptations depend on changes in gene expression (61), we investigated the translated mRNA in NAC D1SPNs and D2SPNs. We focused on relatively stable alterations by isolating cell population-specific translating mRNA one day after the last training session. We used 4 samples per condition, each sample containing the left and right NAc from 3 mice with a proportion of female mice in each sample 0.3-0.5 (**Supplementary Table 1**). Our experiment (see **Figure 1A**) was designed to detect the mRNA footprints of i) the goal-directed component of food-seeking behavior (master vs. yoked), ii) food palatability (yST vs. yHP) and iii) the interaction between food palatability and goal-directed behavior (mST vs. mHP). We retained the genes that exhibited an absolute change greater than 1.5-fold in either direction (absolute log2 fold-change over 0.585), with a p-value adjusted for multiple comparison less than 0.05 (summary in **Table 1**). We previously showed that the genes detected with this approach are predominantly originating from SPNs (46).

**Table 1.**
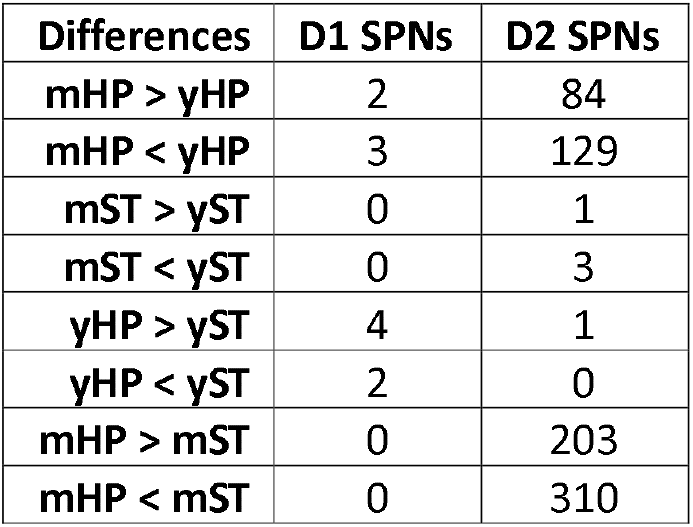
Number of differentially expressed genes.

| Differences | D1 SPNs | D2 SPNs |
| --- | --- | --- |
| mHP > yHP | 2 | 84 |
| mHP < yHP | 3 | 129 |
| mST > yST | 0 | 1 |
| mST < yST | 0 | 3 |
| yHP > yST | 4 | 1 |
| yHP < yST | 2 | 0 |
| mHP > mST | 0 | 203 |
| mHP < mST | 0 | 310 |

We detected almost no differences in D1R-expressing neurons (**Table 1, Supplementary Tables 3-6**). In contrast, operant conditioning for HP food had striking effects in D2R-expressing neurons (**Table 1, Supplementary Table 7**). Although cholinergic interneurons also express D2R, the translating mRNA was mostly originating from D2SPNs as indicated by the low levels of cholinergic interneurons markers including Chat, Slc10a4, Slc18a3, Slc5a7, and Slc17a8 (**Supplementary Table 7**), as previously reported (46). Changes in translated mRNA between mHP and yHP were detected in 213 genes (**Figure 3A, Supplementary Table 7**). Operant conditioning for ST food had few effects on translated mRNA (4 genes, **Table 1, Supplementary Table 8**). The type of food caused almost no differences between yHP and yST mice (1 gene, **Table 1, Supplementary Table 9**). In contrast, modifications were detected between mHP and mST in 513 genes (**Table 1, Figure 3B, Supplementary Table 10**). Thus, most changes in mRNA were observed in D2SPNs and in response to operant training for HP food. The complete results of the comparison between operant conditioning for HP food (mHP vs yHP) and the comparison between conditioning with HP or ST food (mHP vs mST) are highly correlated (R = 0.87, p < 10^−15^, **Figure 3C**). This reflects the predominant effects of conditioning with HP food on gene expression in NAc D2-neurons (**Table 1**). Accordingly, among the 213 mRNAs differentially translated between mHP and yHP, 140 (66%) were also significantly changed between mHP vs mST, all changing in the same direction (**Supplementary Table 11**). The major conclusion of the translatome analyses was that most changes induced by operant training were taking place in D2SPNs and resulted from up or downregulation of translated mRNA in the mHP group as compared to the others.

**Figure 3:**
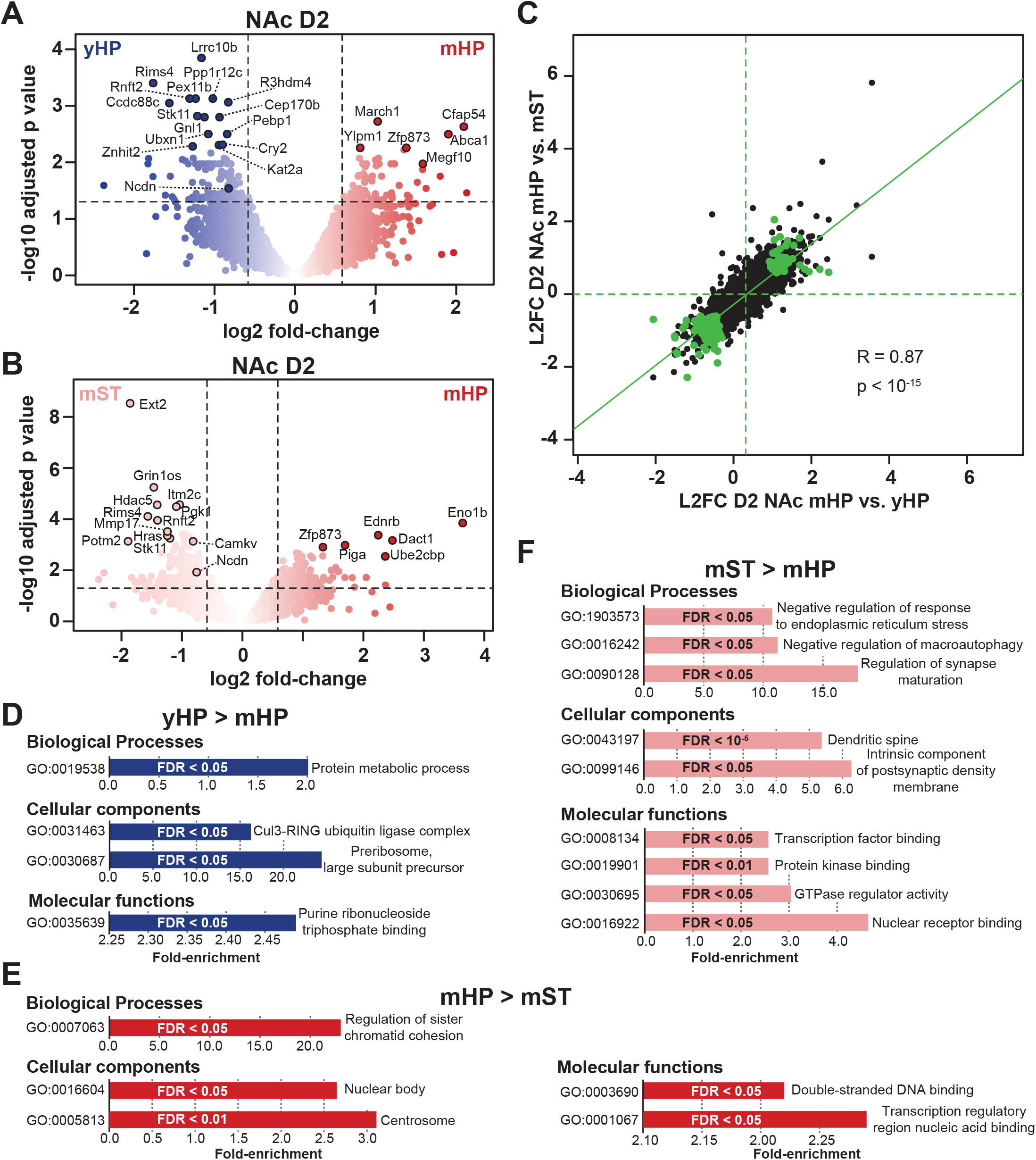
Effects of operant training for standard and highly palatable food on translated mRNA in D2R-expressing neurons of the nucleus accumbens (NAc). Drd2-EGFP/Rpl10a transgenic mice trained as described in Figure 1A were killed 24 h after the last training session and the NAc was rapidly dissected. mRNA was immunopurified (3 mice pooled per sample) and quantified by the TRAP-seq method (**Table 1, Supplementary Tables 7** and **10**). **A**) Volcano plot of genes comparison between yHP and mHP groups in NAc D2R-expressing neurons. Names of main differentially expressed genes are indicated. **B**) Same as in A) for mST vs mHP groups. **C**) Scatter plot of the translated mRNA differences in the NAc D2 SPNs between mHP vs. yHP groups (X axis, see A) and between mHP vs. mST (Y axis). The correlation coefficient was calculated for all genes. mRNAs significantly different in both comparisons are indicated in green. **D**) Gene ontology analysis of genes differentially more expressed in yHP than in mHP in D2R-expressing neurons (**Supplementary Table 12**). Only the main non-redundant GO pathways are indicated. **E-F**) Same as in D) for genes more expressed in mHP than in mST (**E**) or more expressed in mST than in mHP (**F**) (**Supplementary Table 13**). **D-F**, False discovery rate (FDR) values are indicated.

### Pathways and gene interaction clusters affected in NAc D2SPNs by operant conditioning

To evaluate the functional implications of the changes observed in translated mRNAs induced by operant conditioning for HP food in D2SPNs we used several approaches. First, gene ontology analysis (GO) showed that the pathways decreased in D2SPNs of mHP mice as compared to the yHP group were related to ribosomes and ubiquitin ligase (**Figure 3D, Supplementary Table 12**). In the comparison between mHP and mST, GO pathways increased in mHP were related to chromatin and centrosomes, whereas those that were diminished included synaptic and signaling pathways (**Figure 3E and F, Supplementary Table 13**). We then used the network inference performed as previously described (46) to identify gene clusters affected by operant conditioning with HP food. The output was a large connected network with only a few isolated edges (**Figure 4**). The main network presented two super-connected clusters of nodes with degrees over 10 (**Figure 4**). Both clusters comprise mostly genes whose expression was decreased by conditioning. One is enriched in genes coding proteins involved in calcium and cAMP signaling (*Camkv, Mast3, Pde1b, Ppp1r12c, Tesc*) possibly indicating a modulation of signaling, including by dopamine. This cluster also contains GABA receptor subunit delta (*Gabrd*) and alpha3 (*Gabra3*), higher in yHP and mHP, respectively. *Ncdn* that codes for neurochondrin, a.k.a. norbin, is also part of this cluster, decreased in mHP. The second cluster is enriched in proteins involved in protein production, from transcription (*Gtf2f1, Zpf622*) and splicing (*Puf60, Rrp1, Znhit2*) to translation (*Eif2b5, Rplp0, Trmt61a*), folding (*Cct7, Vbp1*) and protein modifications (*Gm16286, Ube3a*). This cluster analysis indicates that beyond synaptic function, conditioning also regulates basic cellular properties and suggests overall adaptation of cellular properties following operant conditioning for HP food.

**Figure 4:**
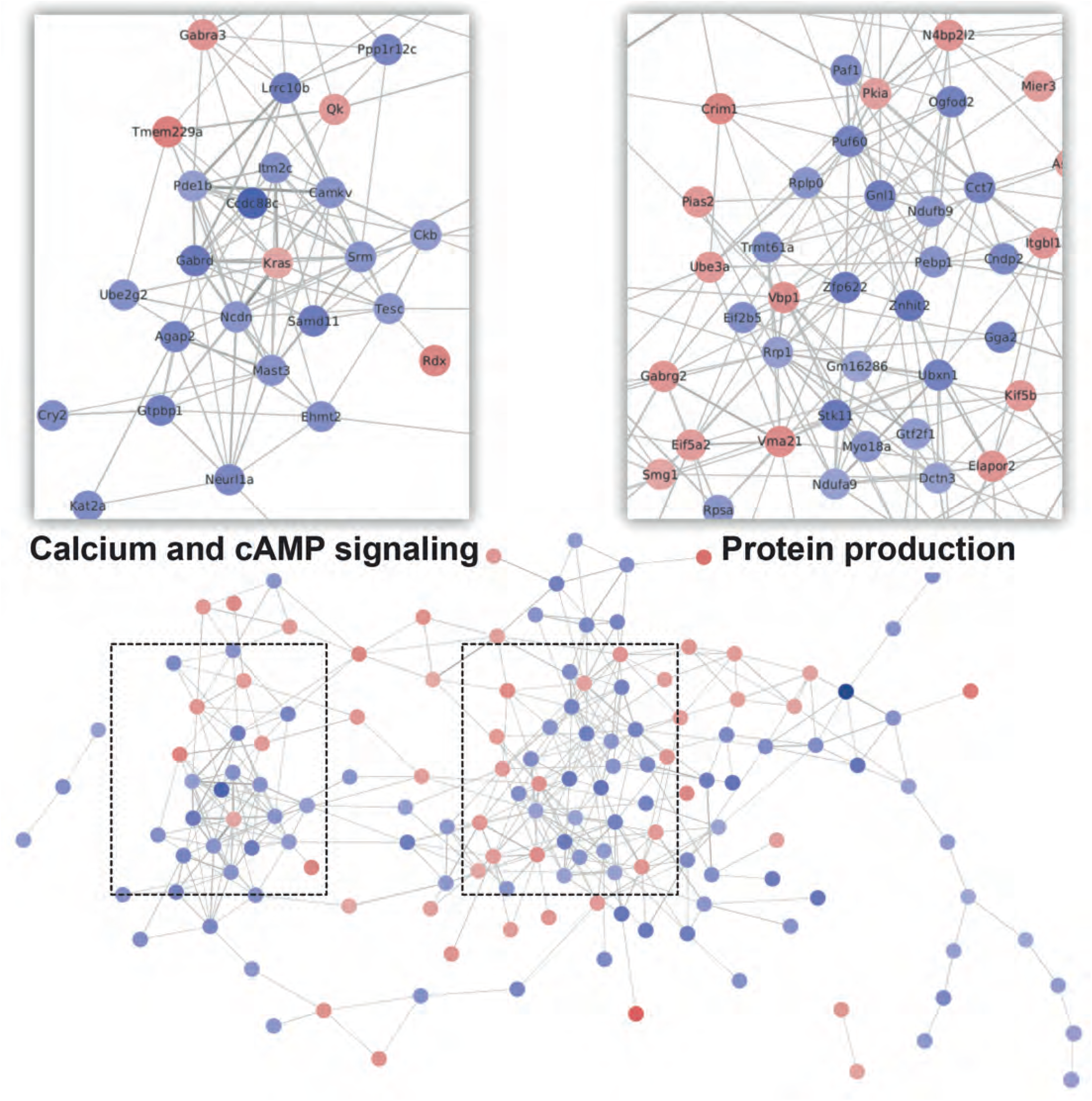
Main clusters of genes with translated mRNA regulated by conditioning for highly palatable food in the D2R-expressing neurons of the nucleus accumbens. Only the top 1% predicted interactions were retained and filtered using the genes with a mean expression greater than 301CPM, differentially expressed by at least 50% (log2 fold change greater than 0.585) at an FDR less than 0.05. Almost all the remaining genes formed a super-cluster. Two over-connected sub-clusters emerged, with degrees over 10. They comprised mostly genes suppressed by highly palatable food, involved either in calcium and cAMP signaling (left) or protein production (right). Nodes are colored with log2 fold changes of expression, blue genes being more expressed in yoked animals and red genes more expressed in master animals. Edge thickness and darkness are proportional to the interaction score.

### Role of Ncdn in motivation for HP food

Having identified the impact of HP food seeking on the translational landscape in dopaminoceptive neurons and their predominance in D2SPNs, we then sought to validate the relevance of our analysis by genetically manipulating one of the genes differentially regulated by conditioning for HP food in this neuronal population. We focused on *Ncdn*, which was highly expressed and downregulated by conditioning for HP food in NAc D2-SPNs, as compared to yHP and mST groups (**Supplementary Table 1**). This gene is of particular interest because it codes for neurochondrin (a.k.a. norbin), which is an important modulator of morphological and synaptic plasticity through the regulation of mGluR5 membrane trafficking (47) and Ca^2+^/calmodulin-dependent protein kinase 2 (CaMK2) activity (62). Moreover, forebrain-specific *Ncdn* KO generates a depressive-like phenotype (47). We hypothesized that deletion of *Ncdn* in neurons could mimic or alter behavioral features of operant responding for HP food. We used previously characterized *Ncdn* conditional KO (*Ncdn*-cKO) mice (47, 63), generated by crossing *Ncdn*^Flox/Flox^ with *Camk2a*-Cre mice and compared them to their *Ncdn*^Flox/Flox^ littermates, which did not express Cre, designated as *Ncdn*^F/F^ mice below. Both *Ncdn*-cKO and *Ncdn*^F/F^ mice displayed higher operant responding when working for HP food than for ST (**Figure 5A, B**, and see **Supplementary Table 2** for all statistical analyses). The response of *Ncdn*-cKO mice for HP food was similar or at some time points higher than that of *Ncdn*^F/F^ mice (**Figure 5A, B**) showing that the mutation did not impair operant learning but appeared to increase it during food-deprived FR1 and FR5 reinforcement schedules. When mice had ad *libitum* access to food, we noted slightly different changes in the rate of pokes depending on the food and genotype. In *Ncdn*^F/F^ mice a decrease in positive pokes was observed for ST food (**Figure 5C**), but not for HP food (**Figure 5D**), as in WT mice (**Figure 1F**). In contrast, a pronounced decrease in nose pokes was observed in *Ncdn*-cKO mice working for HP food (**Figure 4D**). This result suggested that despite their stronger learning phase than *Ncdn*^F/F^ mice (**Figure 5A, B**) the *Ncdn*-cKO mice were less motivated for HP food when they had ad *libitum* access in their home cage. We tested this hypothesis of a reduced motivation of *Ncdn*-cKO mice for HP food using a progressive ratio protocol one day after the conditioning paradigm (**Figure 5E**). The *Ncdn*-cKO mice displayed a lower breaking point than the *Ncdn*^F/F^ mice indicating a decreased motivation (**Figure 5E**).

**Figure 5:**
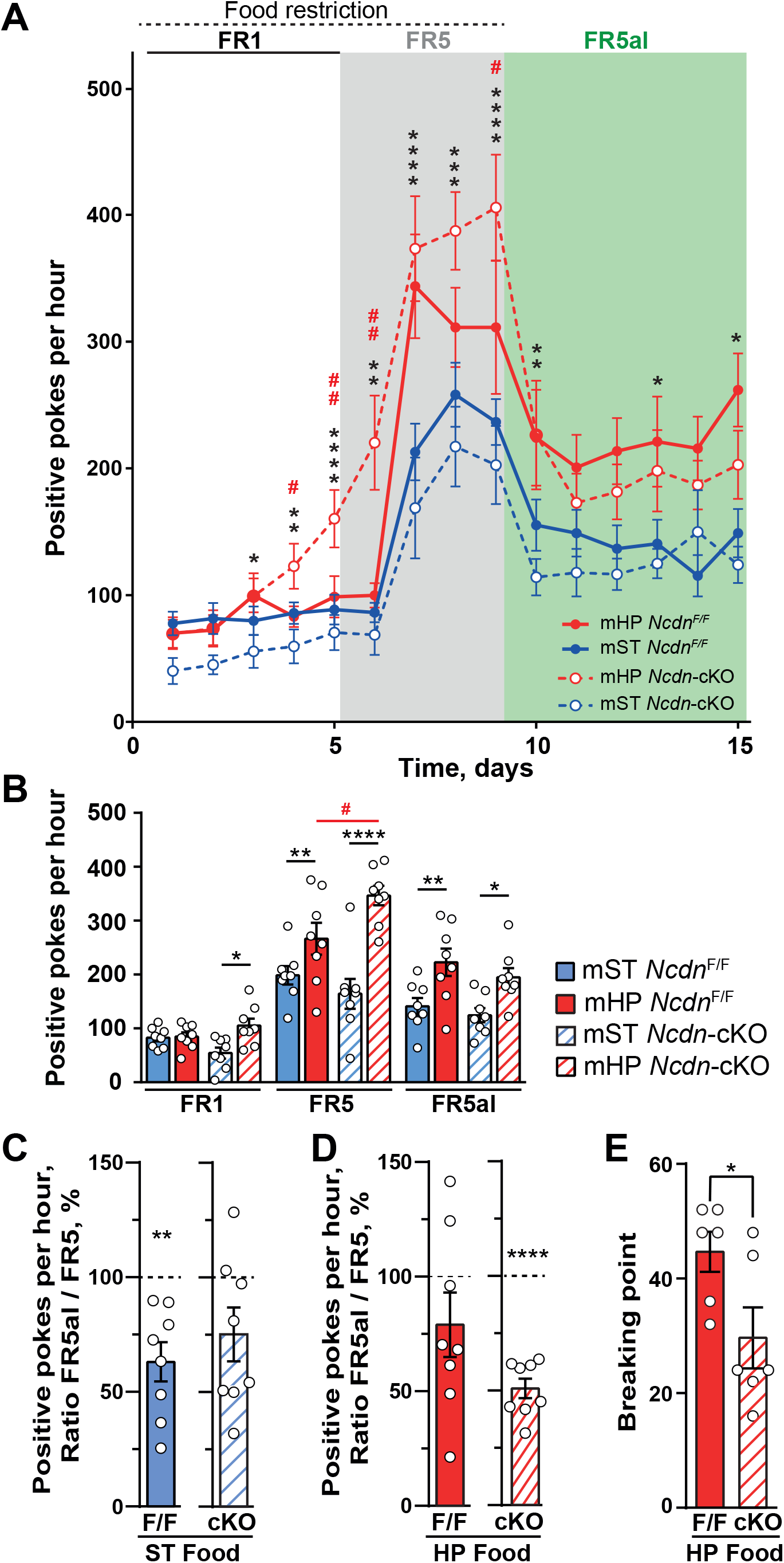
Role of *Ncdn* in satiety-induced devaluation. **A**) Time-course of the number of positive pokes (i.e. in the active hole) during operant training of ***Ncdn*-cKO** and control littermates (***Ncdn***^F/F^) for **ST** or **HP** food (8 mice per group), using the schedule described in Figure 1A. Two-way repeated measure ANOVA for each phase (see **Supplementary Table 2** for complete statistical results). Group effect, FR1, F_(3, 28)_ = 4.6, p<0.01, FR5, F_(3, 28)_ = 11.42, p<10^−4^, FR5al, F_(3, 28)_ = 6.17, p = 0.002. Fisher’s LSD between mST and mHP *Ncdn*-cKO (*) and mHP in *Ncdn*^F/F^ vs *Ncdn-KO* (red #). **B**) Average positive pokes per 1-h session during each period of conditioning, for standard (**mST**) or highly palatable (**mHP**) food in ***Ncdn-cKO*** and ***Ncdn***^F/F^ mice (8 per group, data from Figure 4A). Two-way ANOVA, group (genotype/food) effect, F_(3, 63)_ = 17.5, p < 10^−4^, conditioning phase effect, F_(3, 63)_ = 86.9, p<10^−4^, interaction, F_(6, 63)_ = 3.00, p = 0.012. Fisher’s LSD between mST and mHP *Ncdn-cKO* (*) and mHP in *Ncdn*^F/F^ vs *Ncdn-KO* (red #). **C-D**) In these mice (8 per group) satiety-induced decreased motivation for nose-poking was evaluated as in Figure 1D by making the positive pokes ratio FR5al/FR5. (**C**) for ST food, one sample t test, *Ncdn*^F/F^, 8 mice, t_7_ = 4.33, p = 0.0034, *Ncdn-cKO*, 8 mice, t_7_ = 2.17, p = 0.069. (**D**) for HP food, one sample t test, Ncdn^F/F^, 8 mice, t _7_= 1.50, p = 0.18, Ncdn*-cKO*, 8 mice, t _7_= 11.5, p <10^−4^. **E**) The motivation of Ncdn^F/F^ and Ncdn*-cKO* mice (6 per group) trained for HP food was evaluated in a progressive ratio protocol, in which the number of pokes necessary to obtain the reward is increased after each reward. The breaking point is the highest number of pokes for one pellet achieved by each mouse. Mann and Whitney test, U = 5, p = 0.032. **A, D-E**, * p<0.05, ** p<0.01, **** p<10^−4^.

To determine whether these operant results depended on a change in reward sensitivity, we tested *Ncdn*^F/F^ and *Ncdn*-cKO mice in a free choice paradigm over 43 days, during which mice of either genotype had free access to both ST and HP food. The analysis of cumulative caloric intake showed that in this free choice condition both *Ncdn*^F/F^ and *Ncdn*-cKO mice had a ∼20-fold preference for HP over ST food (**Figure 6A**). However, HP food intake was significantly lower in Ncndn-cKO mice as compared to *Ncdn*^F/F^ controls (**Figure 6A**). *Ncdn*^F/F^ mice in the free choice condition displayed a significant weight gain as compared to another group with access to ST food only (**Figure 5B**). In contrast in *Ncdn*-cKO mice this weight gain in free choice condition as compared to ST only was not observed (). Thus, deletion of *Ncdn*, a gene whose mRNA is less translated in NAc D2-SPNs following operant conditioning for HP food, tends to increase operant behavior during food-deprived conditions, but restores satiety-induced devaluation and decreases motivation for HP food and weight gain in a free choice paradigm. These latter results suggest that decreased expression of *Ncdn* in the NAc could be a compensatory mechanism, counterbalancing the persistent effects of HP food on motivation.

**Figure 6:**
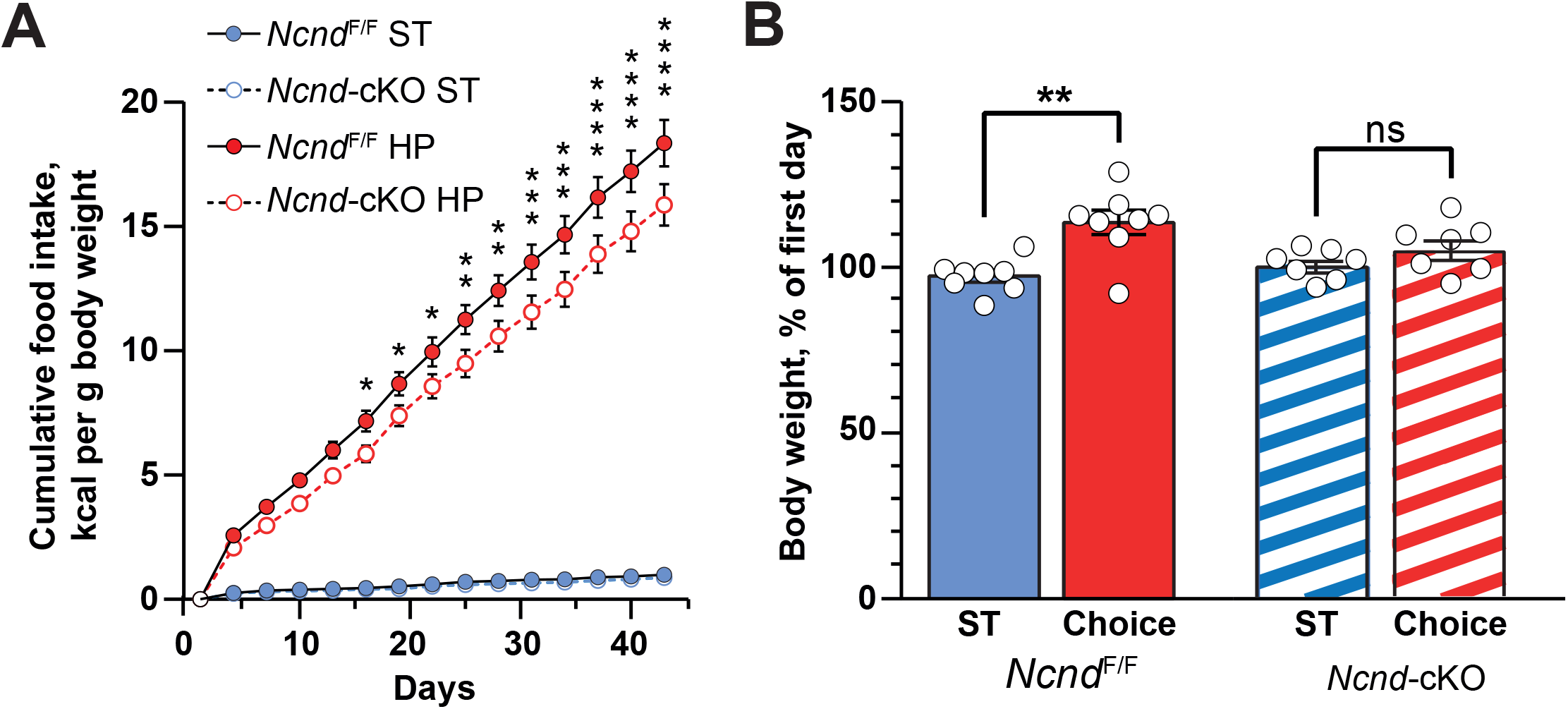
*Ncdn cKO* decreases HP food consumption and weight gain in food free choice conditions. **A**) *Ncdn-cKO* mice consume less HP food than Ncdn^F/F^ littermates in a free choice paradigm. For 6 weeks, mice in their home cage had free access to ST or HP food and the consumption of each type of food was monitored (*Ncdn*^F/F^, n = 8 mice, *Ncdn*-cKO, n = 7 mice). Although mice from both genotypes clearly preferred HP food the Ncdn-cKO animals consumed less HP food than Ncdn^F/F^ controls. Two-way ANOVA, interaction F_(42, 364)_ = 207.3, p<10^−4^, time effect F_(14, 364)_ = 736.7, p<10^−4^, group effect, F = 201.3, p<10^−4^. Post hoc multicomparison Holm-Sidak’s test results are shown between HP food consumption of *Ncdn*-cKO and *Ncdn*^F/F^ mice. Significant differences in consumption of HP food are indicated, * p<0.05, ** p<0.01, *** p<0.001, **** p<10^−4^. **B**) The weight of mice in free choice conditions in A) was compared to the weight of mice of the same genotype with access only to ST food. The weight of *Ncdn*^F/F^ mice in free choice was higher than in ST food-only conditions, whereas this was not the case of *Ncdn*-cKO mice. Kruskal-Wallis test, p = 0.0066, Dunn’s multiple comparisons test, *Ncdn*^F/F^, p = 0.0018, n = 8 mice per group, *Ncdn*-cKO, p = 0.55, n = 7 mice per group.

## DISCUSSION

Highly palatable foods recruit and have the potential to hijack the brain reward system, like drugs of abuse, leading to overeating, satiety-induced desensitization, and craving, thus resulting in weight increase and contributing to increased prevalence of obesity [see (3, 6, 9, 23, 64) for reviews]. The mechanisms of these effects of food are typically investigated in rodent models, using free-access to fat- and/or sugar-rich food. Even though these paradigms mimic in part the “western diet”, they do not disentangle the metabolic and rewarding effects of the food or probe the effects of palatability on goal-directed actions. Here, using operant conditioning with food rewards that only differ by their palatability, HP and ST food, we monitored the effects of palatability on two main components of food-seeking behavior, believed to be altered in pathological feeding behavior, i.e. the incentive component and sensitivity to satiety. As expected, HP food enhanced motivation to exert effort in both food-deprived and ad *libitum* conditions, as described for highly caloric/palatable foods (60). Although both the ST and HP groups significantly decreased responding when switched from restricted to ad *libitum* access to food, the HP group was less sensitive to this satiety-induced devaluation, resembling the accelerated development of habit behavior in animals chronically exposed to high-fat diet (65). This suggests that palatability, independently of caloric intake, could be sufficient to lead to compulsive-like behavior towards food. The reinforcing effect of rewards involve the meso-accumbens dopamine pathway (27, 66). The involvement of the NAc in our experimental conditions is supported by the increase of dendritic spine density in this region after 2 weeks of exposure to highly palatable food, in line with previous reports of longer experiments (60, 67).

Our study identifies long-term alterations in the translatome of dopaminoceptive neurons in the NAc that accompanied HP food-induced long-term behavioral adaptations and persisted at least 24 hours after the last operant conditioning session. Translated mRNAs were mostly altered in response to operant conditioning for HP food, and these alterations were virtually restricted to D2SPNs. Despite the central role of D1SPNs in early food-induced long-term behavioral adaptations (68, 69) we found few changes in the translatome of these neurons at the end of the operant protocol. The persistent changes in D2SPNs align with indirect observations that point to the role of these neurons in addiction, feeding disorders and obesity (5, 70, 71). Interestingly, striatal hypofunction and increased body mass index are more frequent in individuals harboring the ANKK1 TaqIA A1 allele, which is associated with decreased striatal availability of D2R (72-74). Data in rodents also support a role of D2R in the development of some features of obesity (2, 39). Recent findings in mice highlight a particular sensitivity of D2SPNs to circulating lipids (75, 76), and point to a role of D2R and D2SPNs in the regulation of energy output (77, 78). Interestingly GO pathways and gene networks downregulated in NAc D2SPNs of mHP mice included signaling and synaptic functions, while those upregulated were related to DNA and transcription, possibly indicating long-term adaptive modification of these neurons.

To start testing the functional importance of the genes disclosed by our analysis, we focused on *Ncdn* a highly expressed gene, involved in synaptic function and downregulated in the D2SPNs of the NAc. *Ncdn* neuronal deletion had restricted consequences on HP food-related behavior. *Ncdn*-cKO mice showed an increased initial operant response for HP food, suggesting that the basic mechanisms of reward-induced procedural learning were not impaired. However, the mutant mice displayed satiety-induced devaluation for HP food and decreased motivation for HP food in a progressive ratio task performed when mice were fed *ad libitum*. The same *Ncdn*-cKO mutation was reported to generate a depressive-like phenotype (47) and a decreased preference for sucrose (63). In our experiments *Ncdn*-cKO preferred HP food to ST food, and readily learned to work for it. Yet this preference was slightly blunted and did not result in the weight gain observed in control mice. These results suggest a partial alteration of motivation in *Ncdn*-mutant mice, which will be interesting to explore in the context of major depression, a condition in which the reward system is altered (79, 80).

In conclusion, our study provides the first description of the footprint of HP food-operant conditioning on the translating mRNA landscape in dopaminoceptive neurons of the NAc. It underlines the involvement of D2SPNs and identifies networks of implicated genes. One of the downregulated genes, *Ncdn*, may play a compensatory role and oppose the conditioning-induced adaptation of feeding behavior and persistent motivational drive to eat HP food. Further exploration of *Ncdn* actions is warranted to explore novel approaches to counteract overeating and its contribution to obesity.

## Acknowledgments

The authors thank Pierre Trifilieff for helpful suggestions. The present work was supported in part by Inserm and Sorbonne University, by ERC-2009AdG_20090506, *Fondation pour la Recherche Médicale*, (FRM) DEQ20081213971, and ANR-16-CE16-0018 Epitraces to J.A.G., NIH grants DA018343 and DA040454 to A.C.N. Y.N. was supported in part by the Fyssen Foundation and by a Uehara Memorial Foundation fellowship. BdP was supported by the FRM, FDT201805005390.

## Author contributions

JAG, EM, JPR conceived and JAG and JPR supervised the project. EM, JPR, AG, DH, EV, MM, NG, JAG designed the experiments. EM, AG, YN, BdP, AP, MB, JPR performed experiments. EM, JPR, AG, ACN, EV, DH, and JAG analyzed data, LT and NG, performed, analyzed and interpreted bioinformatics analyses, EM, AG, LT, YN, LG, ACN, EV, DH, NG, PG, MF, JPR, and JAG discussed the data and provided input and corrections to the manuscript. EM, NG, JPR, JAG wrote the manuscript. All the authors but PG approved the final version of the manuscript.

