## Supplementary Method for "Operant training for highly palatable food alters translating mRNA in nucleus accumbens D2 neurons and reveals a modulatory role of *Neurochondrin*"

**Supplementary Methods**

**Operant conditioning experiments**

Operant conditioning experiments were carried out in Med Associate operant chambers (model ENV-307A-CT) in an isolation box equipped with a fan. Each box included a grid floor (model EVV-414), 2 nose-poke holes (one randomly selected as active and the other as inactive), one house light, a food dispenser and a food magazine between the 2 nose-poke holes. The beginning of the session was concomitant with the fan activation and the turning on of the house light for 3 sec, and a pellet delivery. The session ended after 60 min or after 100 pellets had been delivered. Seven days before the beginning of the experiment, mice were individually housed and maintained in an environment with controlled temperature and humidity with a 12:12-h reversed light dark cycle. All the experiments were carried out during the dark phase of the dark/light cycle. Five days before the start of conditioning all the mice (including the yoked) were food-restricted to maintain their weight stable, until the 9^th^ operant training session in order to facilitate the acquisition of the task, and then had *ad libitum* food access. During the operant conditioning sessions animals were presented with either 20 mg dustless precision ST pellets (TestDiet 5UTM #1811143) or HP isocaloric pellets (TestDiet 5UTL #1811223). The ST food was similar to that used to maintain the mice (TestDiet Purina 5053) in composition and taste (3.30 kcal g-1, 24.1 % protein, 10.4 % fat, 65.5 % carbohydrates, wt/wt). The HP pellets were similar in calorie content to the standard diet (3.48 kcal/g) but contained a higher level of sucrose among the carbohydrates (49% wt/wt) and had a chocolate flavor. The progressive ratio (PR) schedule lasted for 1 h and respected the following series: 1, 2, 3, 4, 5, 6, 7, 8, 10, 12, 14, 16, 18, 20, 22, 24, 28, 32, 36, 40, 44, 48, 52, 56, 64, 72, 80, 88, 96, 104, 112, 120, 128, 136, 144, 152, 160, 168, 176, 184, 192, 200, 208, 216, 224, 232, 240, 248, 256, 264, 272, 280, 288, 296, 304, 312, 320, 328, 336, 344, 352, 360, 368, 376, 384, 392, 400, 408, 416, 424, 432, 440, 448, 456, 464.

***mRNA extraction for TRAP-Seq***

Cell type-specific ribosome-bound mRNA was purified as described (1, 2) with some changes. TRAP transgenic mice were sacrificed by decapitation. The brain was quickly dissected out, placed in cold buffer and then in an ice-cold brain form to cut thick slices from which the nucleus accumbens (NAc) was punched out using an ice-cold stainless steel cannula. Samples containing tissue pieces from 1-3 mice (final proportion of males 0.46-0.50 in each cell population, **Supplementary Table 1**) were homogenized in 1 mL of lysis buffer (20 mM HEPES KOH, pH 7.4, 5 mM MgCl_2_, 150 mM KCl, 0.5 mM dithiothreitol, 100 µg mL^-1^ cycloheximide (Sigma, #C7698-1g), supplemented with protease (Roche, #11836170001) and RNAse inhibitors (Ambion, #AM2694, Fisher, #PR-N2515) with successively loose and tight glass-glass 2-mL Dounce homogenizers. Homogenates were centrifuged at 2,000 x g, at 4°C, for 10 min. The supernatant was separated from cell debris, and supplemented with NP-40 (EDM Biosciences, 10 µL mL^-1^) and 1,2-dihexanoyl-sn-glycero-3-phosphocholine (DHPC, Avanti Polar lipids, 30 mM, final concentrations). After mixing and a 5-min incubation on ice, the lysate was cleared for 10 min at 20,000 x g. A mixture of streptavidin-coated magnetic beads was incubated 35 min at room temperature with biotinylated protein L and then 1 h with EGFP antibody and then added to the supernatant and incubated overnight at 4°C with gentle end-over-end rotation. Beads were collected with a magnetic rack, washed 4 times with high-salt buffer (20 mM HEPES-KOH, pH 7.4, 5 mM MgCl_2_, 350 mM KCl, 10 µL mL^-1^ NP-40) and immediately placed in “RTL plus” buffer (Qiagen). mRNA was purified using RNeasy Plus Micro Kit (Qiagen) and in-column DNAse digestion. RNA integrity was evaluated using a Bioanalyzer with an RNA Pico Chip (Agilent), and the quantity of RNA was measured by fluorimetry using the Quant-IT Ribogreen kit.

***Libraries and sequencing***

Five ng of RNA were used for reverse-transcription, performed with the Ovation RNA-Seq System V2 (Nugen). cDNA was quantified by fluorimetry, using the Quant-iT Picogreen reagent, and ultra-sonicated using a Covaris S2 sonicator (duty cycle 10 %, intensity 5, 100 cycles per burst, 5 min). Two hundred ng of sonicated cDNA were used for library construction with the Illumina TruSeq RNA sample prep kit, starting at the End-Repair step, and following the manufacturer’s instructions. The libraries were quantified with the Bioanalyzer high-sensitivity DNA kit, multiplexed and sequenced on an Illumina HiSeq 2500 instrument. At least 20 million 50-bp paired-end reads were collected for each sample.

***Bioinformatics analysis***

The quality of the raw data was assessed using FastQC (3) for common issues including low quality of base calling, presence of adaptors among the sequenced reads or any other overrepresented sequences, and abnormal per base nucleotide percentage. The different libraries were then mapped to the *Mus musculus* genome GRCm38 (UCSC mm10) using HISAT2 (4). Depending on the sample, between 37 and 62 million reads were mapped. After RNA-Seq Quality Control, reads were quantified using the RNA-Seq pipeline of SeqMonk (5) and, for each gene product, counts of all reads in all exons were exported with the corresponding gene annotations. Gene products from sex chromosomes were not included in the study. Sequencing data has been deposited in NCBI's Gene Expression Omnibus and are accessible through GEO Series accession number GSE137153 (<https://www.ncbi.nlm.nih.gov/geo/query/acc.cgi?acc=GSE137153>) with the accession numbers indicated in **Supplementary Table 1**. Differential expression was performed with DESeq2 package (6) v1.18 with the option betaPrior set to TRUE. Significance was set at an adjusted p-value of 0.01 and more stringent thresholds were then added to focus on differences most likely to be biologically important (Padj < 10^-3^, L2FC > 1). Except for differential expression analysis, the raw count files were processed to generate counts per million reads (CPM). BaseMeans are means of normalized counts of all samples within a group, normalized for sequencing depth. Data were then normalized with DESeq2’s rlog function.

***Network inference***

Following the conclusions of the DREAM5 challenge’s analysis (7), we used a combination of methods based on different algorithms to infer regulatory networks. We combined the results of CLR (8), a mutual-information-based approach providing undirected edges, and GENIE3 (9), a tree-based regression approach providing directed edges. Both methods were the best performers in their category at DREAM5. Both tools were applied using gene expression from all samples, using filtered CPM as described above. Because CLR only provides undirected edges while GENIE3 provides directed ones, the results of CLR were all mirrored with the same score on edges in both directions. Only edges with a positive score in GENIE3’s results were used. The edges present in both CLR and GENIE3 results were then ranked according to the product of CLR and GENIE3’s scores. This only retains edges that have either an extremely high score with one method, or consistent scores with both methods. Subnetworks were extracted using gene lists as seeds, retaining only the first neighbors with scores above a threshold. Visualization and analysis of the resulting networks was done using Cytoscape (10). The R script is available in ref (11).

***Spine analysis***

Fresh brain hemispheres were processed following the Golgi-Cox method as described (12). Mouse brain hemispheres were incubated in the dark for 21 days in filtered dye solution (10 g L^-1^ K_2_Cr_2_O_7_, 10 g L^-1^ HgCl_2_, and 8 g L^-1^ K_2_CrO_4_). The tissue was then washed 3 x 2 min in water and 30 min in 90% EtOH (v/v). Two hundred-µm sections were cut in 70% EtOH on a vibratome (Leica) and washed in water for 5 min. Next, they were reduced in 16% ammonia solution for 1 h before washing in water for 2 min and fixation in 10 g L^-1^ Na_2_S_2_O_3_ for 7 min. After a 2-min final wash in water, sections were mounted on superfrost coverslips, dehydrated for 3 min in 50 %, then 70, 80 and 100% EtOH, incubated for 2 x 5 min in a 2:1 isopropanol:EtOH mixture, followed by 1 x 5 min in pure isopropanol and 2 x 5 min in xylol. Bright-field images of Golgi-impregnated *stratum radiatum* dendrites from nucleus accumbens principal neurons were captured with a DM6000 (Leica) light microscope equipped with a CCD camera (×100 oil objective). Only fully impregnated pyramidal neurons with their soma found entirely within the thickness of the section were used. Image *z* stacks were taken every 0.2 μm and at 1,024 × 1,024 pixel resolution, yielding an image with pixel dimensions of 49.25 × 49.25 μm. *Z*-stacks were deconvolved using the Huygens software (Scientific volume imaging, Hilversum, Netherlands), to improve voxel resolution and reduce optical aberration along the *z*-axis. Segments of proximal apical dendrites were selected for the analysis of spine density and spine morphology according to the following criteria: (a) segments with no overlap with other branches that would obscure visualization of spines and (b) segments either “parallel” to or “at acute angles” relative to the coronal surface of the section to avoid ambiguous identification of spines. Only spines arising from the lateral surfaces of the dendrites were included in the study; spines located on the top or bottom of the dendrite surface were ignored. Given that spine density increases as a function of the distance from the soma, reaching a plateau 45 μm away from the soma, we selected dendritic segments of basal dendrites 45 μm away from the cell body. The total number of spines was obtained using the cell counter tool in the ImageJ software. At least 40 dendrites per group from at least eight mice per group were counted. The imaging and analysis were done by different investigators who were both blind to the mouse group.
